# Expression analysis of photosynthesis-related genes in albino *Artocarpus heterophyllus* seedlings leaves

**DOI:** 10.1101/2020.10.20.346833

**Authors:** Zeping Cai, Junna Dong, Xi Zhang, Qian Qu, Fanhua Wu, Lu Cao, Shidong Li, Zixuan Wang, Dan Zhou, Jiajia Luo, Xudong Yu

**Author notes:** Corresponding author, (XY). These authors contributed equally to this work (Zeping Cai, Junna Dong, Xi Zhang).

## Abstract

Albino *Artocarpus heterophyllus* Seedlings (AAS) were found in the preliminary investigation by our group and were used as materials for researching. The phenotype of AAS leaves were observed and measured. In parallel, the photosynthetic physiological parameters were determined under different photosynthetically active radiations (PAR). The results suggested that the length, width, area and thickness of AAS leaves were less than normal seedings. Likewise, the net photosynthetic rate, intercellular CO_2_ concentration, stomatal conductance and transpiration rate of AAS leaves were not susceptible to PAR in contrast to normal individuals. Furthermore, the transcriptome sequencing technology was performed to clarify the expression of genes related to photosynthesis. It is as expected that numerous down-regulated genes were found in the synthesis of photosynthetic pigments, as well as the pathways of photosynthesis - antenna proteins, photoreaction and carbon fixation reaction of AAS leaves. Compared to other albino plants, AAS have a longer life span and more stable phenotypic traits with larger leaves, which could provide ideal materials for investigating photosynthesis of woody plants.

## Introduction

Photosynthesis is the basis of the development of green plants, supplying the necessary energy for virtually all life on earth. It plays an important role in maintaining the relative stability of oxygen and carbon dioxide ratios in the atmosphere [1–3].

The capture, absorption, transmission and transformation of light energy in photosynthesis are mainly undertaken by photosynthetic pigments [4]. Lacking of photosynthetic pigments may result several variations in leaf color, such as albinism, yellowing and other phenomena [5]. Albino seedlings rely on the nutrients in the seeds to maintain growth, and these seedings would die once the nutrients are exhausted. So far, the researches on albino plants have been mainly carried on *Oryza sativa* [6], *Arabidopsis thaliana* [7], *Zea mays* [8], *Nicotiana tabacum* [9] and so on. The seeds of these plants are generally smaller with less nutritious and can only live a short life after they germinated.

There are two stages in photosynthesis: photoreaction and carbon-fixation reaction. In the photoreaction stage, H2O can be oxidized and decomposed by light energy to synthesize high-energy compounds (ATP and NADPH). While in carbon-fixation reaction, the ATP and NADPH were used to drive the synthesis of organics by a series of enzymatic reactions [10]. Albino seedlings are ideal materials to study photosynthesis [11]. Previous researches on photosynthesis have emphasized on the herbaceous plants mostly whereas only a few reports were found in the woody plants, such as *Camellia sinensis* [12] and *Citrus reticulata* [13].

*Artocarpus heterophyllus* is a perennial evergreen tree with high economic value [14]. During the preliminary investigation, our research group found albino *Artocarpus heterophyllus* seedlings (AAS) whose mother plants had recessive heritable albinism [15]. Compared with other plants, *A. heterophyllus* seeds are larger and able to store more nutrients. Therefore, AAS have a longer life span lacking of energy provided by photosynthesis. Moreover, AAS, whose leaves are larger, show innate advantages for researching photosynthesis. In this study, AAS leaves were used for morphological structure observation and photosynthetic physiological parameters determination. And their leaves were also regarded as the samples for transcriptome sequencing to analyze the expression of related genes. Above all, the research could provide materials for investigating photosynthesis in woody plants.

## Materials and methods

### Acquisition of experimental materials

The materials in this study were from Danzhou City, Hainan Province (109°29’11’’E, 19°30’29’’N), and their mother plant was a heterozygote *A. heterophyllus* tree with recessive heritable albinism (S1 Fig). The seeds were stripped from mature fruits. Plump seeds were selected and planted in pots containing adequate nutrient soil under the ample light environment. After their germination, the albino *Artocarpus heterophyllus* seedlings (AAS) in the progeny seedlings were used as the experimental materials and we selected the normal individuals used as control check (CK).

### Methods

#### Measurement of Length, width and area of leaves

The development profiles of the third leaf of AAS and CK were observed within 20 days from the moment when the seedings germinated. The length, width and area of the leaves obtained from at least six plants were measured.

#### Leaves fixation and paraffin sections

The mature leaves of AAS and CK were taken to make paraffin sections following the method described by Jagadish et al. [16]. After that, the photos photographed by LEICA DFC295 optical microscope were used to measure the leaf thickness, the upper and lower epidermis thickness, the palisade and sponge tissue thickness and so on.

#### Measurement of photosynthetic physiological parameters

The mature leaves of three AAS and three CK plants were used for physiological detections with the LI-6400XT portable photosynthesis system [17]. Respiratory rate, net photosynthetic rate, intercellular CO_2_ concentration, stomatal conductance and transpiration rate were determined under PAR of 1600, 1400, 1200, 1000, 800, 600, 400, 200 and 0 μmol·m^−2^·s^−1^.

#### Determination of chlorophyll and carotenoids content

The photosynthetic pigments in mature leaves of AAS and CK were extracted and their content was determined according to the method of Arnon [18]. Absorbance (A) was determined at the wavelength of 663 nm (A_663_), 646 nm (A_646_) and 470 nm (A_470_) with 95% ethanol as the control check, which was repeated 3 times. The content of chlorophyll a, chlorophyll b, total chlorophyll and carotenoids were respectively calculated as the following formulas:

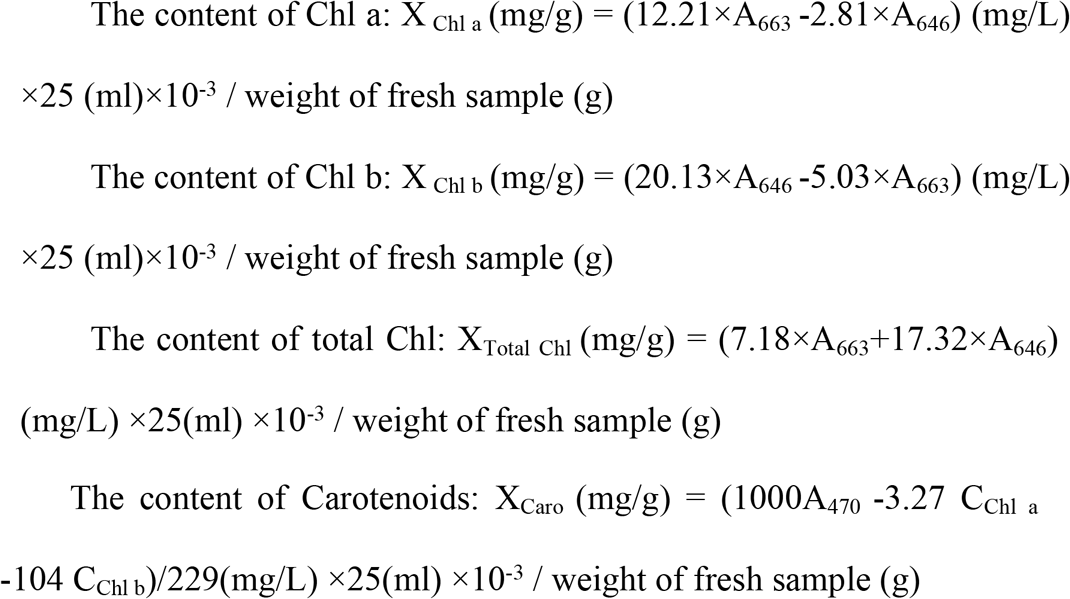

#### Data measurement and statistical analysis

The length, width and area of AAS and CK leaves as well as the thickness of the epidermis, sponge tissue and palisade tissue of the leaves were all measured by Image J, for which six or more repetitions were set. The mean and standard deviation were calculated, and statistically significant differences were analyzed by U-test between the CK and AAS data.

#### RNA extraction, cDNA library construction and transcriptome sequencing

The mature leaves of AAS and CK were put into the mortar with liquid nitrogen, ground into powder and transferred into EP tubes containing lysate liquid. Total RNA was extracted by CTAB [19]. Afterwards, polyA mRNAs were highly enriched with magnetic oligo(dT) beads. The enriched mRNAs were fragmented by fragmentation buffer, and the fragmented mRNAs were used as a template for reverse transcription to compound double-stranded DNA after synthesis of cDNA two-strand. Sequencing connectors were added to amplify the target fragments by PCR. The single-stranded DNA generated by thermal denaturation of PCR products was cyclized to obtain single stranded circular DNA library. BGISEQ-500 high-throughput sequencing platform was used for sequencing. The construction of cDNA library and sequencing were entrusted to Beijing Genomics Institute (BGI).

#### Gene assembly, KEGG annotation and analysis

The raw reads were filtered by removing the reads with low quality, connector pollution and the content of unknown base N more than 5%. The filtered clean reads (PRJNA661997) were compared to the full-length transcriptome reference gene sequences of *A. heterophyllus* (PRJNA579273) in the SRA database using Bowtie2. *De novo* assembly was subsequently carried out to obtain the final Unigenes. All Unigenes with a length of more than 200 nt were subjected to BLAST search against the database of KEGG for annotation (E-value < 1e-5).

The Unigenes with sequence similarity greater than 70% were termed into the same Cluster and named after ‘CL’. Unigenes of related pathways were selected, the expression levels of each Unigene in the Cluster were summed up. The Clusters and Unigenes with average expression levels less than 0.5 were removed, while the rest of them with significant difference (t<0.5) were selected by T-test to draw the heatmaps.

#### Quantitative real-Time PCR (qRT-PCR) analysis

Fifteen genes were selected for qRT-PCR analysis [20]. The primers were designed by Primer Express Software v2.0 of ABI. In addition, the reactions were carried out on ABI ViiA 7 PCR amplifier, with three parallel experiments conducted for each sample. Relative quantification was performed by 2^−ΔΔCt^ method with *CL11555(AhUBIQUITIN)* as an internal reference gene.

## Results

### Measurement of the length, width and area of AAS leaves

The whole leaves of AAS are white. With the increase of growth aging, the quicker increases were observed of the length, width and area in CK leaves than AAS in the first 17 days. AAS leaves reached the maximum in the length, width and area which showed significant differences from CK on the 17th day. They were 41.07 mm, 21.47 mm and 704.53 mm^2^, Which value 43.13%, 36.05% and 20.81% of those in CK (95.23 mm、 59.54 mm and 3385.49 mm^2^), respectively. The larger aspect ratio was observed in AAS leaves in contrast to CK, and significant differences were shown from the 12th day (Fig 1c-f). The daily growths of the length, width and area of AAS leaves were all less compared with CK during the whole growth period. The length and width of the leaves increased rapidly in the first 9 days, but tended to be gentle in 9-17 days. While in the days after the 17th day, the length, width and area showed negative growth, which is due to the premature atrophy of AAS leaves (Fig 1f-h).

**Fig 1.**
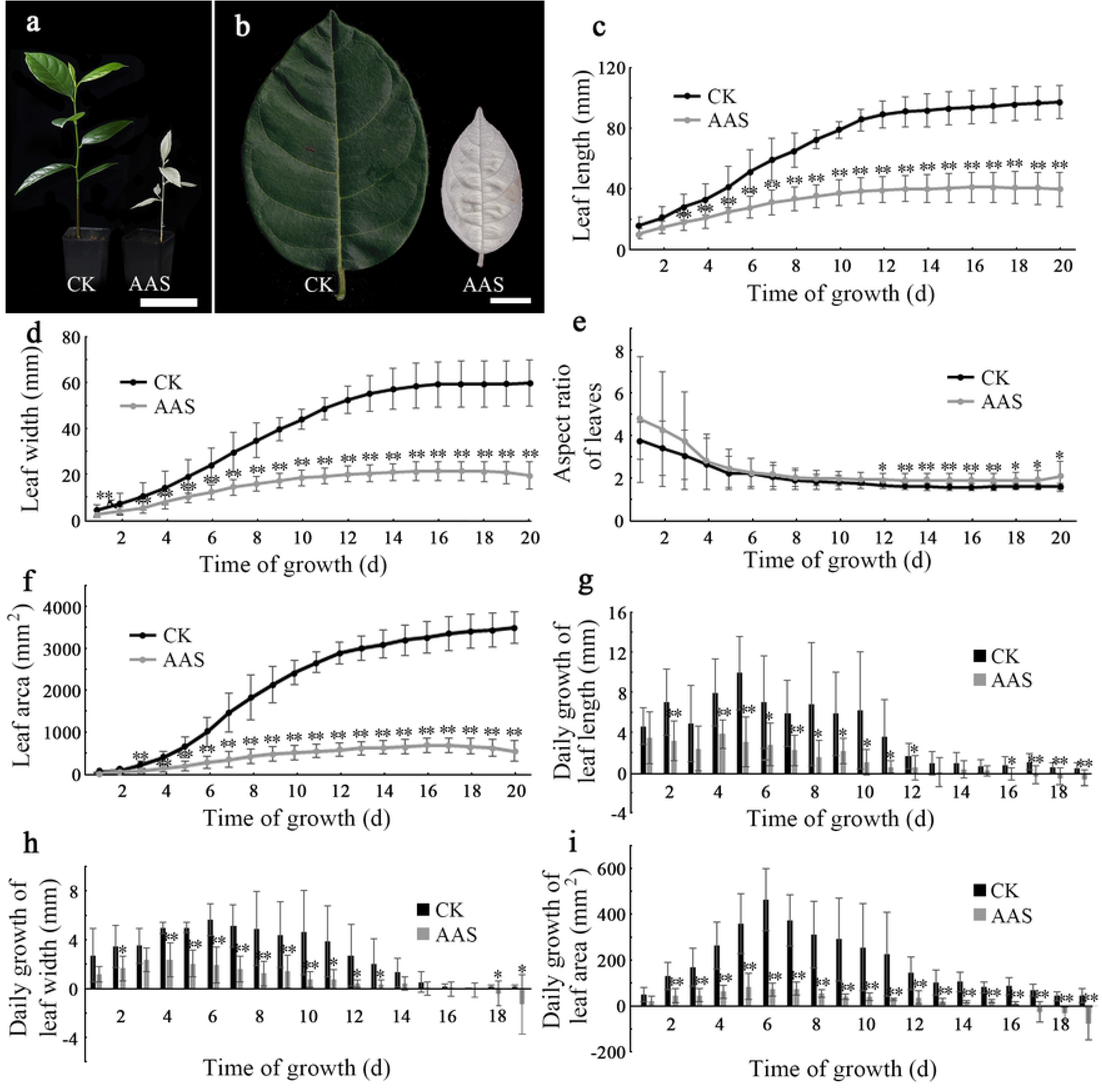
Analysis of leaf length, width and area in AAS. (a-b): Phenotype of plants (a) and leaves (b); (c-f): Daily changes of leaf length (c), leaf width (d), leaf aspect ratio (e) and leaf area (f); (g-i): Daily growth of leaf length (g), leaf width (h) and leaf area (i). The scale of a is 10 cm; that of b is 1 cm. *: Significant difference, *P*<0.05; **: Extremely significant difference, *P*<0.01; CK: control check; AAS: Albino *Artocarpus heterophyllu*s Seedlings, the same below.

### Analysis of the leaf epidermis and mesophyll of AAS

The thicknesses of the upper and lower epidermis of AAS leaves were both less than those in CK, which were 14.68 μm and 10.97 μm, accounting for 86.40% and 67.93% of those in CK (16.99 μm and 16.15 μm), respectively. Both values had significant differences from those in CK (Fig 2b-c). The thicknesses of palisade tissue and spongy tissue of AAS were 31.34 μm and 152.79 μm, which were 79.99% and 59.73% of those in CK (39.18 μm and 255.82 μm), respectively. While no significant differences were observed in the cell width of palisade tissue and spongy tissue between AAS and CK. The thickness of AAS leaves was less than that in CK, which was 215.15 μm and 64.25% of that in CK, with significant difference compared with CK (Fig 2g).

**Fig 2.**
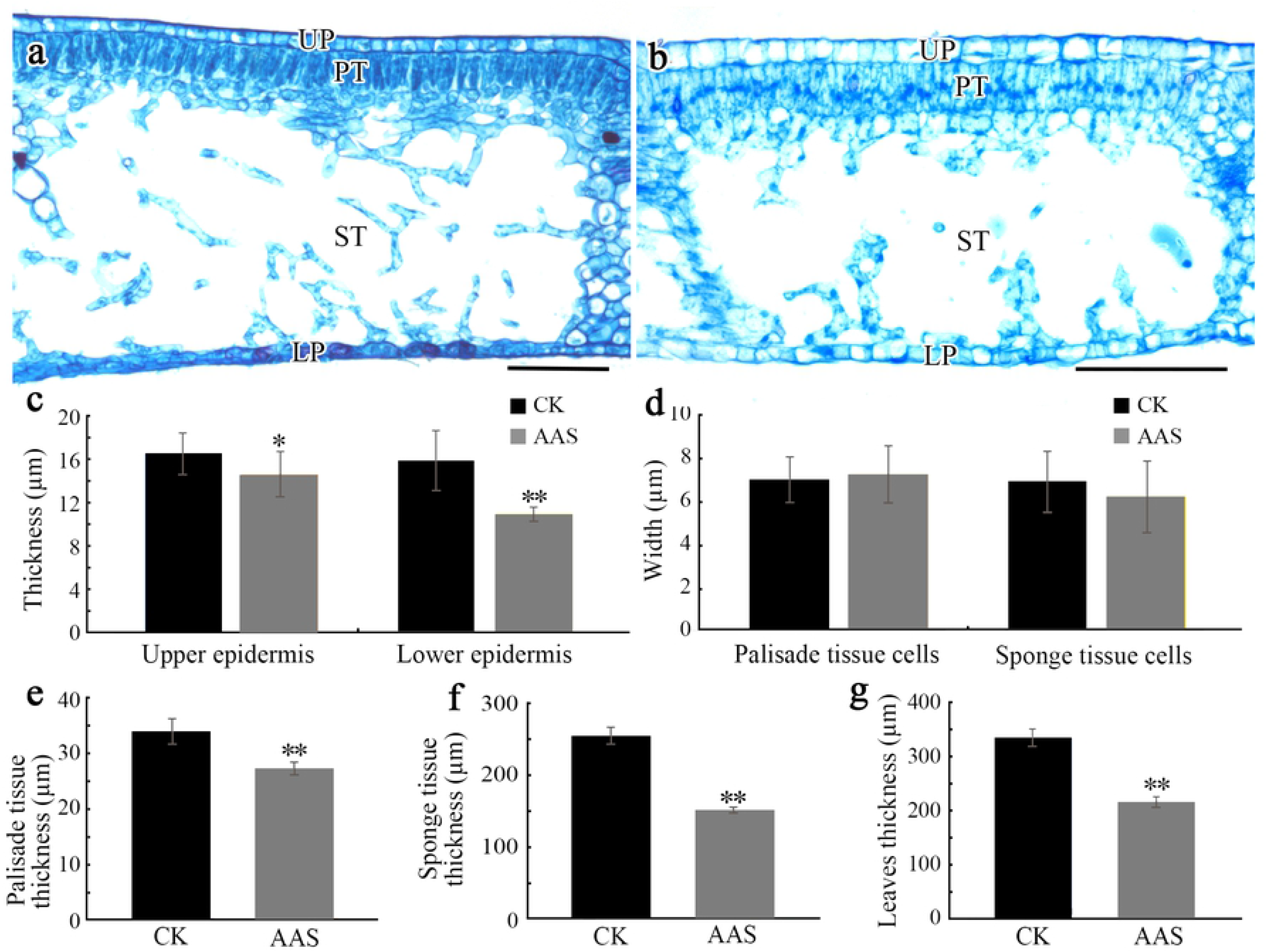
Analysis of leaf epidermis and mesophyll in AAS. (a-b): Leaf cross sections of CK (a) and AAS (b); (c): Thickness of upper and lower epidermis; (d): Cell width of palisade tissue and sponge tissue; (e-g): Thickness of palisade tissue (e), sponge tissue (f) and leaves (g); The scale is 100 μm. UE: Upper epidermis; LE: Lower epidermis; PT: Palisade tissue; ST: Sponge tissue.

### Photosynthetic physiological analysis of AAS leaves

The contents of Chlorophyll a, Chlorophyll b, total Chlorophyll and Carotenoids in AAS leaves were 0.013 mg·g^−1^, 0.014 mg·g^−1^, 0.027 mg·g^−1^ and 0.0077 mg·g^−1^, which decreased down to 1.07%, 1.67%, 1.32% and 1.28% of CK levels (1.21 mg·g^−1^, 0.84 mg·g^−1^, 2.05 mg·g^−1^ and 0.60 mg·g^−1^), respectively (Fig 3a). The respiratory rate of AAS leaves was 0.916 μmol·m^−2^·s^−1^, which was 2.18 times of that in CK with statistical significance (Fig 3b). From 0 to 1600 μmol·m^−2^·s^−1^ PAR, only slight fluctuations were observed in the net photosynthetic rate of AAS leaves. And the similar trend was also shown in the intercellular CO_2_ concentration, the stomatal conductance and the transpiration rate of AAS leaves as expected (Fig 3c-f). Increasing the PAR from 0 to 600 μmol·m^−2^·s^−1^ resulted in an enhance in the net photosynthetic rate, while conducted a decline simultaneously in the intercellular CO_2_ concentration of CK leaves. With further increases in PAR, the net photosynthetic rate of CK leaves essentially saturated, and little fluctuations were observed in the intercellular CO_2_ concentration (Fig 3c-d). With the increase of PAR, the stomatal conductance of CK leaves increased gradually. The observation is consistent with the trend of transpiration rate in CK which varied from that in AAS significantly (Fig 3e-f). These results clearly showed that AAS leaves were not susceptible to PAR compared with CK.

**Fig 3.**
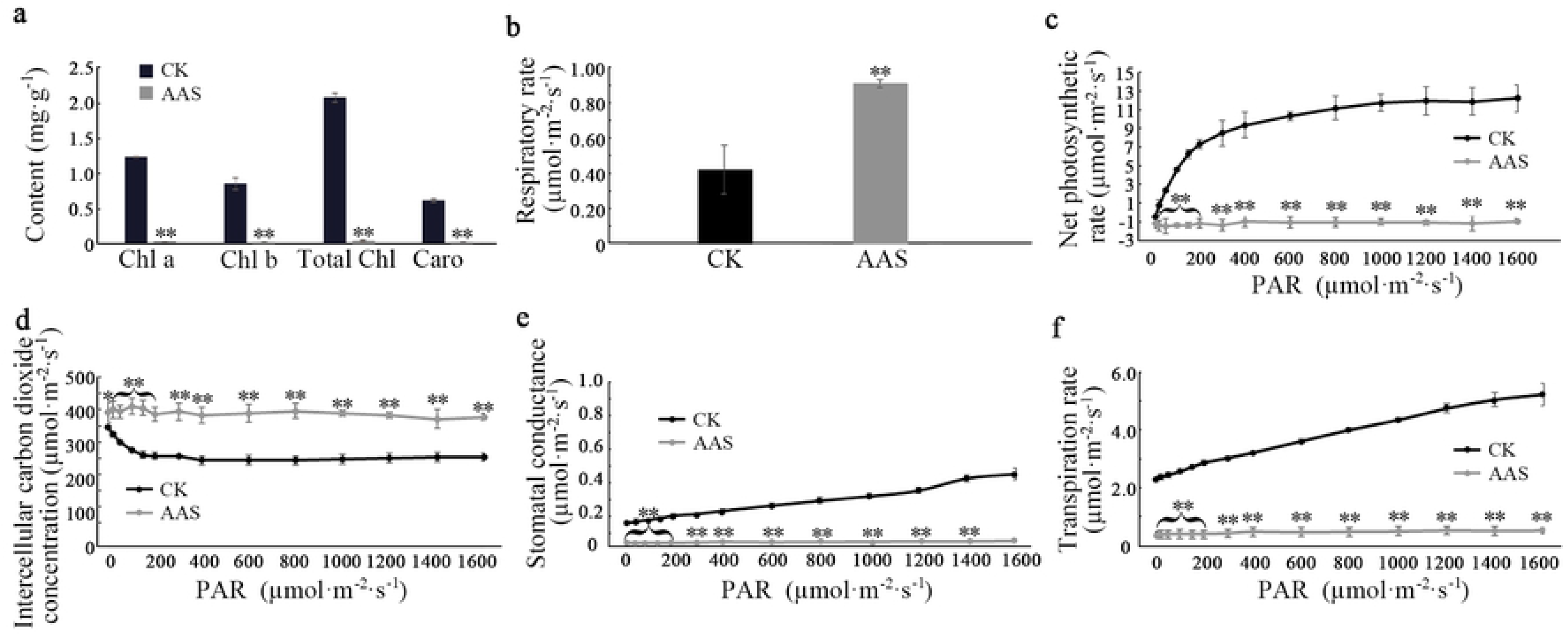
Analysis of leaf photosynthetic physiological in AAS. (a): The contents of chlorophyll a, chlorophyll b, total chlorophyll and carotenoids; (b): Respiratory rate of leaves in CK and AAS; (c-f): The changes of net photosynthetic rate (c), intercellular carbon dioxide concentration (d), stomatal conductance (e), transpiration rate (f) with the increase of PAR of leaves in CK and AAS. Chl a: chlorophyll a; Chl b: chlorophyll b; Total Chl: total chlorophyll; Caro: carotenoids.

### Sequencing quality assessment

The mRNAs of the 6 samples of AAS and CK leaves were sequenced. The results showed: a total of 437 million raw reads were generated. Afterwards, 414.63 million clean reads were obtained after filtration. Ultimately, 148 440 Unigenes were obtained by subsequently assembling, clustering and removing redundancy. The length of Unigenes ranged from 200 nt to more than 3000 nt. The total length, average length and N50 length of Unigenes were 241 624 864 nt, 1 627 nt and 2 429 nt, respectively. Furthermore, the GC content was 41.25% (Table 1).

**Table 1.** Summary of transcriptome data for *A. heterophyllus*.

|  | Number | Percentage |
| --- | --- | --- |
| Raw reads | 437 million |  |
| Clean reads | 414.63 million |  |
| All-Unigene | 148,440 |  |
| Unigenes (200–500 nt) | 37,046 | 24.96% |
| Unigenes (500–1500 nt) | 41,957 | 28.27% |
| Unigenes (1500–3000 nt) | 49,010 | 33.02% |
| Unigenes ($\geq 3000$ nt) | 20,427 | 13.76% |
| All-Unigene Length (nt) | 241,624,864 |  |
| Mean length (nt) | 1,627 |  |
| N50 length (nt) | 2,429 |  |
| GC |  | 41.25% |

Table 1. Summary of transcriptome data for *A. heterophyllus*.
|  | Number | Percentage |
| --- | --- | --- |
| Raw reads | 437million |  |
| Clean reads | 414.63million |  |
| All-unigene | 148,440 |  |
| Unigenes (200–500 nt) | 37,046 | 24.96% |
| Unigenes (500–1500 nt) | 41,957 | 28.27% |
| Unigenes (1500–3000 nt) | 49,010 | 33.02% |
| Unigenes ($\geq 3000$ nt) | 20,427 | 13.76% |
| All-unigene Length (bp) | 241,624,864 |  |
| Mean length (bp) | 1,627 |  |
| N50 length (bp) | 2,429 |  |
| GC |  | 41.25% |

Data from six sequencing samples were used for Cluster-sample and Pearson correlation coefficient analysis. The results displayed that three biological replicates of CK and AAS were clustered into two branches (S2 Fig). Moreover, the Pearson correlation coefficient among samples of each group were all greater than 0.86 (Fig 4a). The results of principal component analysis indicated that CK and AAS were separated on the first principal component analysis (PCA 1) (Fig 4b). In conclusion, the 6 sequencing samples had high quality and reliable sequencing data.

**Fig 4.**
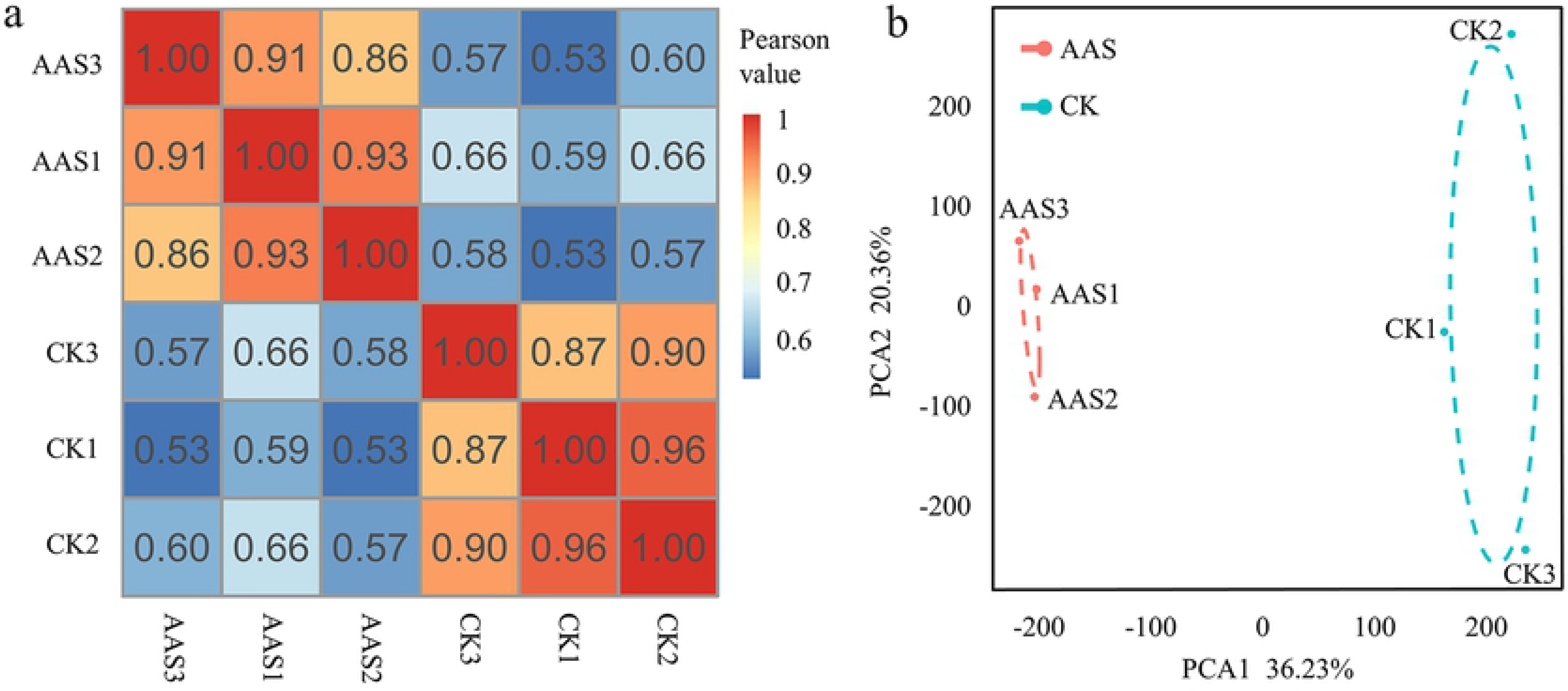
Analysis of Pearson correlation coefficient and principal component. (a): Pearson correlation coefficient analysis was performed among 6 samples using RNA-seq expression data; (b): Principal component analysis (PCA) was performed on 6 samples from two groups using RNA-seq expression data. The diagram shows the separation of each group on the PCA1 and the PCA2.

### Gene expression analysis of chlorophyll and carotenoid synthases

Two categories of photosynthetic pigments are involved in higher plants: chlorophyll and carotenoids.

Chlorophyll a and chlorophyll b were synthesized by a series of enzyme catalytic reactions starting with L-glutamyl-tRNA. Gene expression analyses showed that Glutamyl tRNA reductase gene (*HemA*), Glutamate-1-semialdehyde-2, 1-aminomutase gene (*HemL*), Porphobilinogen synthase gene (*HemB*), Porphobilinogen deaminase gene (*HemC*), Uroporphyrinogen III decarboxylase gene (*HemE*), Magnesium protoporphyrin IX methyltransferase gene (*CHLM*), 3,8-divinyl protochlorophyllide a 8-vinyl reductase gene (*DVR*) and Protochlorophyllide oxidoreductase gene (*POR*) in the process of compounding chlorophyll a and chlorophyll b were all significantly down-regulated. Among these genes, *HemA* and *DVR* were the key enzyme genes (S1 Table). The enzyme genes involved in 10 steps were down-regulated out of the 18 steps of chlorophyll biosynthesis in AAS (Fig 5).

**Fig 5.**
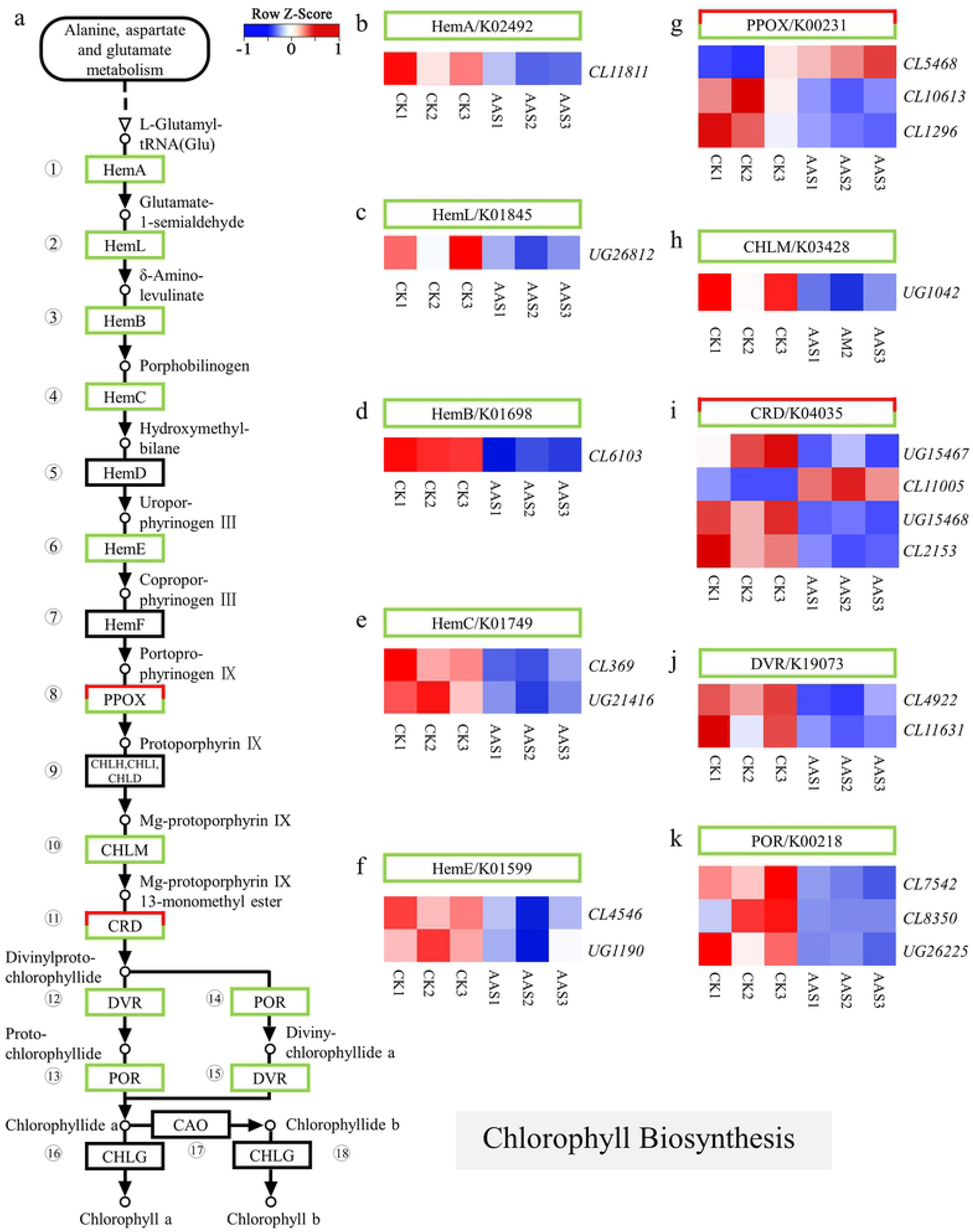
Chlorophyll biosynthesis pathway and expression heatmaps of enzyme genes. (a): Chlorophyll biosynthesis pathway and the expression of related genes; The red / Green / black boxes indicated that the enzyme gene expression was up regulated / down regulated / no significant differences in AAS leaves. (b-k): Heatmaps of enzyme genes *HemA, HemL, HemB, HemC, HemE, PPOX, CHLM, CRD, DVR* and *POR* expression in 6 samples. In this study, only genes with significant difference in expression (*P* < 0.05) were used to map heatmaps, Z-score was used to normalize data, the same below.

It was also found that levels of three key enzyme genes expression of phytoene synthase gene (*PSY*), phytoene desaturase gene (*PDS*), and zeta - carotene desaturase gene (*ZDS*) relative to the synthesis of lycopene from geranylgeranyl diphosphate (GGPP) were lower than CK. The following γ-carotene, δ-carotene, α-carotene, β-carotene and ε-carotene were generated by lycopene respectively, under the effect of the Lycopene β-cyclase (LCYb) and Lycopene ε-cyclase (LCYe). Then, lutein, zeaxanthin and violaxanthin were produced by the catalysis of Carotenoid ε-cyclohydroxylase (LUT1), β-carotene 3-hydroxylase (crtZ) and Violaxanthin de-epoxidase (VDE). Therein, *LUT1* was up-regulated whereas the genes of *LCYb, LCYe* and *VDE* showed significantly down-regulated (S2 Table) in AAS. The enzyme genes associated with 10 steps were down-regulated out of the 16 steps of carotenoids biosynthesis in AAS (Fig 6).

**Fig 6.**
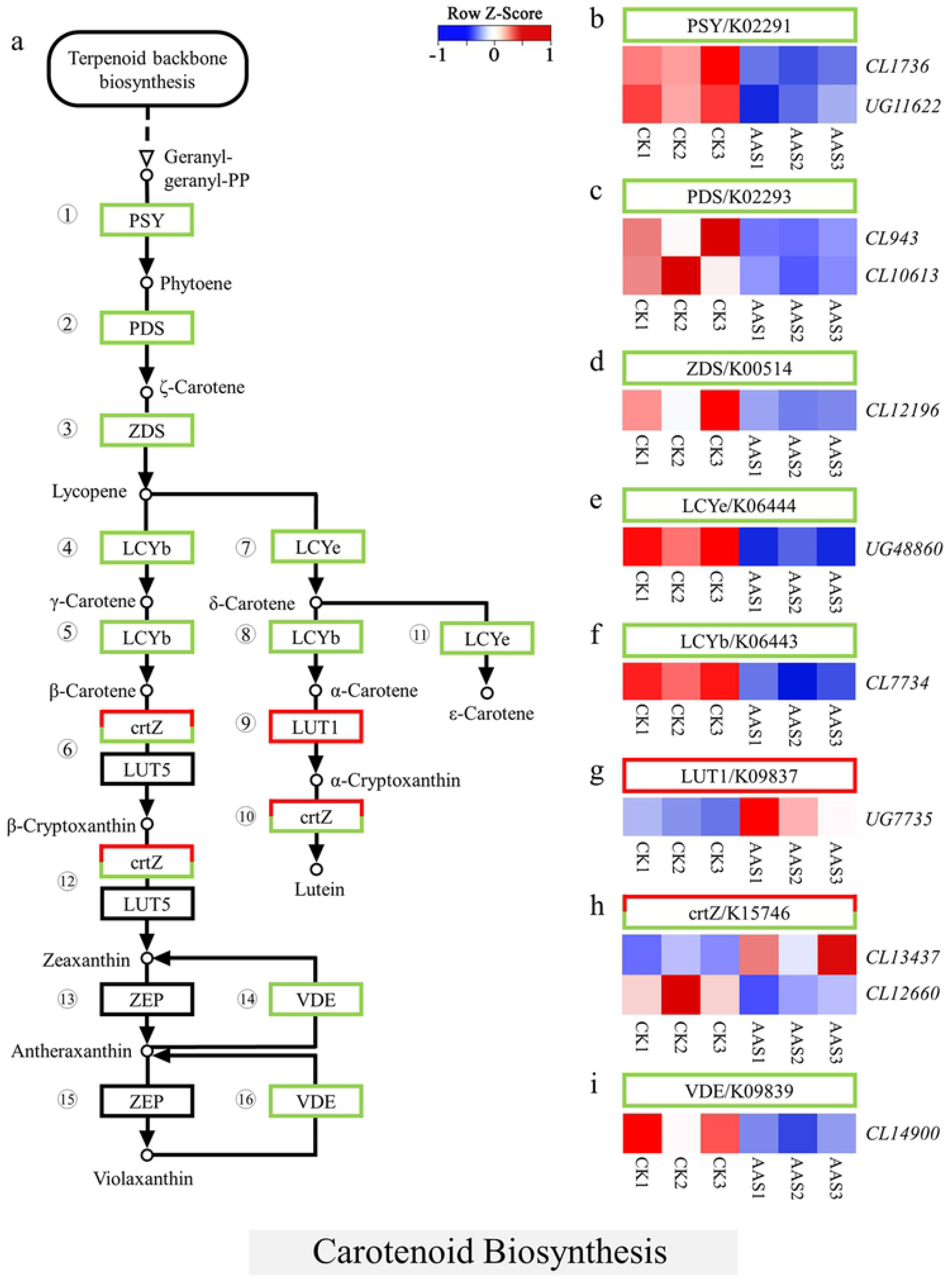
Carotenoid biosynthesis pathway and expression heatmaps of enzyme genes. (a): Carotenoid biosynthesis pathway and the expression of related genes; (b-i): Heatmaps of enzyme genes *PSY, PDS, ZDS, LCYe, LCYb, LUT1, crtZ* and *VDE* expression in 6 samples.

### Expression analysis of related genes in photosynthesis-antenna proteins and photoreaction

In the photosynthesis - antenna proteins pathway, Chl a/b-binding protein complexes (LHC) both in photosystem I (PS I) and photosystem II (PS II) were known to capture light energy and quickly deliver excitation energy to the reaction center. The genes encoding the subunits of LHC were significantly down-regulated in AAS in contrast to CK, except for genes that encoded Lhcb1 and Lhcb7 (Fig 7, S3 Table).

**Fig 7.**
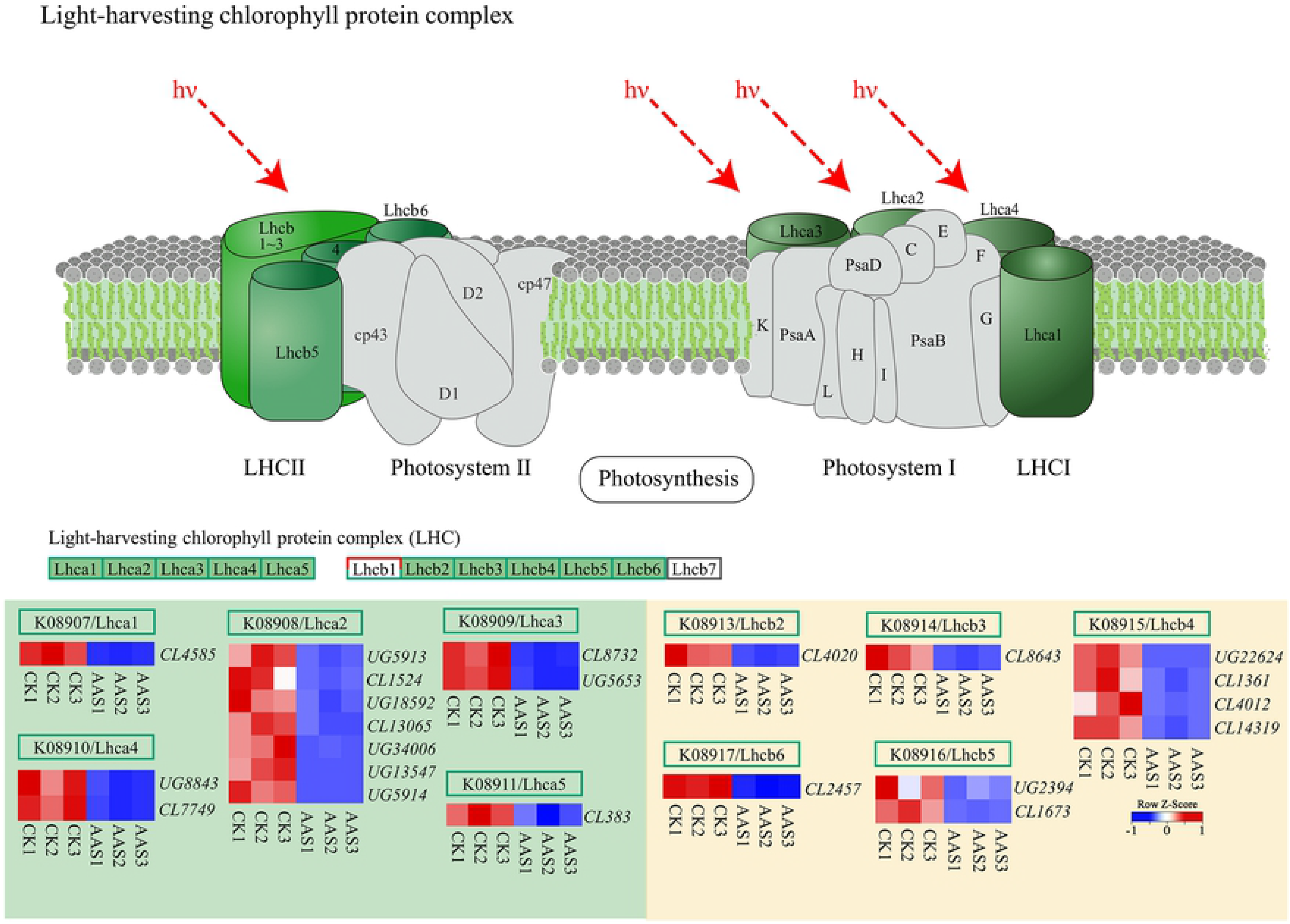
Photosynthesis-antenna proteins pathway and heatmaps of related genes expression. LHC I: the light-harvesting chlorophyll protein complex associated with photosystem I; LHC II: the light-harvesting chlorophyll protein complex associated with photosystem II.

Photosystem I (PS I) and photosystem II (PS II) in photoreaction pathway were mainly in charge of the absorption, transfer, conversion of light energy and the photolysis of H_2_O. The genes coding subunits PsaA、 PsaB、 PsaD、 PsaE、 PsaF、 PsaG、 PsaH、 PsaK、 PsaL、 PsaN and PsaO in PS I, were down-regulated in AAS. Similarly, the coding genes of relevant subunits in PS II, namely, PsbB、 PsbC、 PsbO、 PsbP、 PsbQ、 PsbR 、 PsbS 、 PsbW 、 PsbY and Psb27, were also down-regulated. Besides, the coding genes of subunit PetC in cytochrome b6/f complex (Cyt b6/f), which connects PSII and PSI, showed down-regulation as well. The genes coding constituent subunits beta (β), alpha (α), gamma (γ), delta (δ) and c of adenosine triphosphate Synthetase complex (ATP Synthetase complex), which use proton dynamic potentials formed by cytochrome b6/f complex to drive ATP synthesis, also showed significant down-regulation. In addition, photosynthetic electron transport on thylakoid membrane decompose H_2_O and drove electrons to reduce NADP^+^ by using light energy. During this process, the coding genes of relevant subunits, namely, PetE, PetH and PetJ, were all down-regulated. (Fig 8, S4 Table).

**Fig 8.**
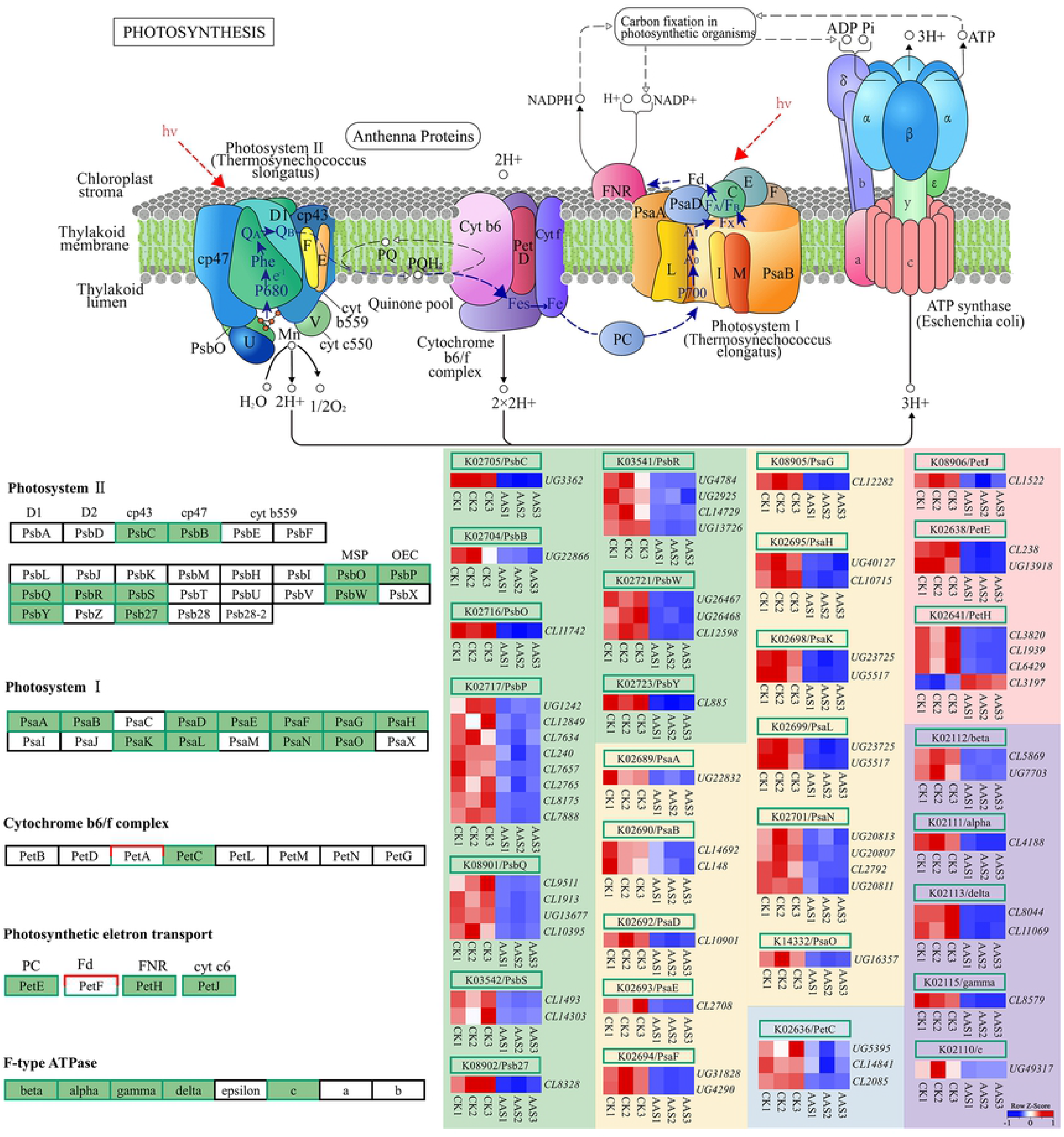
Photoreaction pathway and heatmaps of related genes expression.

### Expression analysis of related genes in carbon-fixation reaction

Calvin cycle is the most common carbon-fixation reaction, including three stages: carboxylation, reduction and regeneration. Primarily, in the carboxylation phase, CO_2_ is attached to Ribulose-1,5-bisphosphate (RuBP) to yield Glycerate 3-phosphate (3-PGA) by the enzyme ribulose-1, 5-biphosphate carboxylase (Rubisco/RBCL). The expression of gene *RBCL* was no significant difference in contrast to CK. In the reduction phase, 3-PGA further yielded Glyceraldehyde-3P (GAP) consuming ATP and NADPH, as activated by Phosphoglycerate kinase (PGK) and catalyzed by Glyceraldehyde 3-phosphate dehydrogenase (GAPDH/GAPA). During this process, gene *GAPDH/GAPA* showed up-regulated significantly. Ultimately, the CO_2_ acceptor RuBP was regenerated by GAP, catalyzed by several enzymes during the regenerative phase. Among them, Fructose 1,6-bisphosphatase gene (*FBP*), Transketolase A, B gene (*TKTA, TKTB*), Phosphoribulokinase gene (*PRK*) were all down-regulated. (S5 Table). The gene expression in 5 steps out of 14 steps in this process were lower than CK obviously (Fig 9).

**Fig 9.**
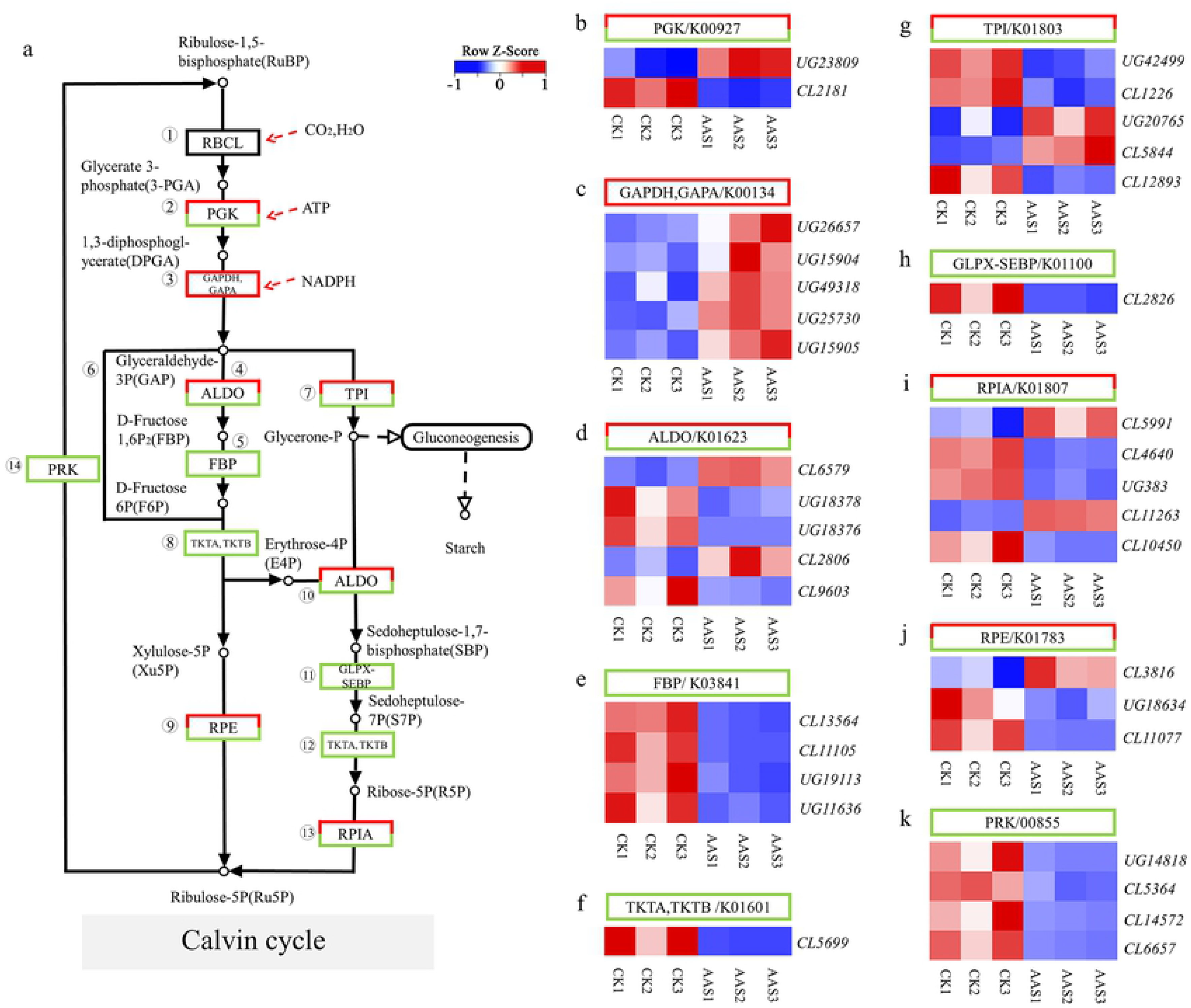
Calvin cycle and enzyme gene expression in carbon fixation reaction. (a): Calvin cycle pathway and the expression of related genes; (b-k): Heatmaps of enzyme genes *PSY*, *PDS, ZDS, LCYe, LCYb, LUT1, crtZ* and *VDE* expression in 6 samples.

### Quantitative real-time PCR (qRT-PCR) analysis

In order to validate the accuracy of RNA-Seq data, we selected 15 genes for quantitative real-time PCR (qRT-PCR) analysis. The 15 selected genes included 5 genes involved in photosynthesis - antenna proteins, 3 genes involved in PS II, 2 genes involved in PS I, 1 gene involved in cytochrome b6/f complex, 2 genes involved in Photosynthetic electron transmission and 2 genes involved in F-type ATPase. The statistical results showed that 15 genes showed the same expression trend as that of RNA-Seq data (Fig 10), indicating the reliability of RNA-Seq sequencing data in this study (S6 Table).

**Fig 10.**
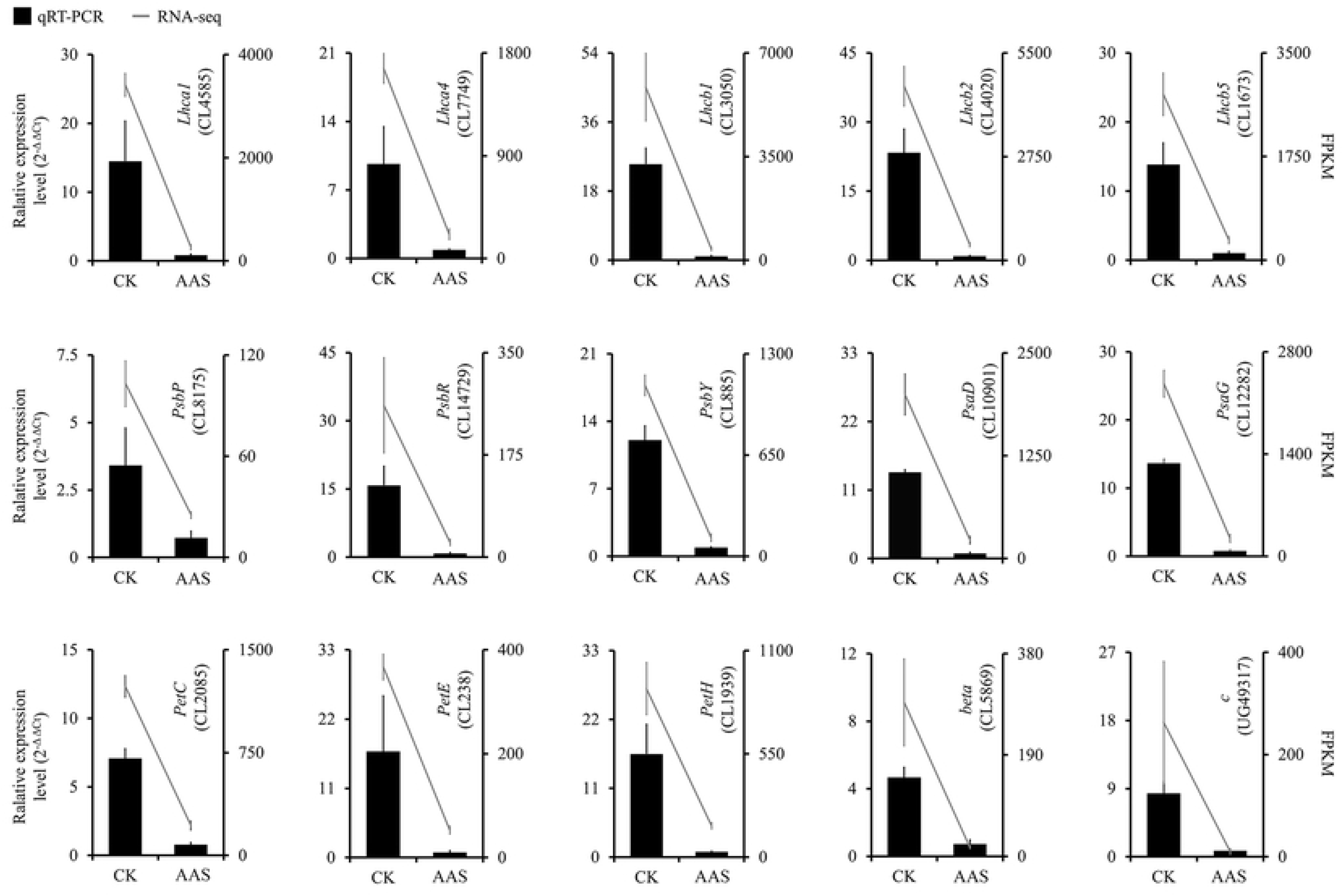
Quantitative real-time PCR (qRT-PCR) analysis of photosynthesis-related genes

## Discussion

Albino plants lacking photosynthetic pigments are ideal experimental materials for studying chloroplast development and photosynthetic mechanism [21–22]. Albino herbaceous plants such as *Oryza sativa* and *Solanum lycopersicum* have been used as breeding materials to cultivate new varieties [23–24]. The albino woody plant, Anji Baicha, has high amino acid content in its leaves and shows the better flavor and quality [25]. Therefore, albino plants are of great significance in scientific research, breeding and quality improvement.

Qin et al. [26] found that the expression levels of *ZDS 、 PSY* and other genes were significantly decreased in *Arabidopsis thaliana* mutant *pds3*, which is due to *PDS*, a gene involved in carotenoids biosynthesis, had mutated. The synthesis of carotenoids was obstructed, which conducted the inhibition in chlorophyll synthesis pathway. The chlorophyll content was extremely low, resulting the plant completely albino. Fang et al. [27] also found that mutating of genes *OsPDS, OsZDS* and *OsLCY* involved in carotenoids synthetase caused the inhibition of carotenoids synthesis in *Oryza sativa*, which can also lead to an extremely low content of chlorophyll in plants. In this paper, the extremely low chlorophyll content in AAS may also be attributed to the inhibition of carotenoids synthesis owing to the down-regulation of *PDS, ZDS, LCY* and other genes (S7 Table).

Wu et al. [28] found that *OsCHLG* mutated and the activity of its coding enzyme decreased in the chlorophyll deficient mutant of *Oryza Sativa*, which led to the inhibition of chlorophyll synthesis. They also found that the expression of genes coding the light-harvesting Chl a/b-binding protein in PSII was also severely inhibited. Chu et al. [29] found that the chlorophyll content decreased in the chlorophyll deficient mutant of *Brassica napus*, and the expression levels of the genes coding photosynthesis related proteins were declined. Moreover, the photosynthetic capacity was also impaired. These findings are consistent with the results in our study that the expression of some genes encoding light-harvesting Chl a/b-binding protein (LHC), photosystem I, photosystem II and photoelectron transport related proteins were all down-regulated in AAS, which implied that the down regulation of the genes encoding related proteins in photosynthesis may be responsible for chlorophyll deficiency.

Photosynthesis is an ancient and complex evolutionary process, and also one of the widely concerned biological frontiers. In this experiment, the expression of genes related to AAS photosynthesis was analyzed, which can provide materials for the investigation of photosynthesis in woody plants.

## Acknowledgments

We want to thank all the participants in this study.

## Supporting information

**S1 Fig. Growth environment of CK and AAS**

**S2 Fig. Cluster-sample analysis of CK and AAS**

**S1 Table. Gene expression levels involved in Chlorophyll Biosynthesis**

**S2 Table. Gene expression levels involved in Carotenoid Biosynthesis**

**S3 Table. Gene expression levels involved in Photosynthesis - antenna Proteins pathway**

**S4 Table. Gene expression levels involved in Photoreaction pathway**

**S5 Table. Gene expression levels involved in Carbon-Fixation pathway**

**S6 Table. Primers sequence information for qRT-PCR analysis**

**S7 Table. Detailed summary of gene expression**

